# Opening of a Cryptic Pocket in β-lactamase Increases Penicillinase Activity

**DOI:** 10.1101/2021.04.14.439842

**Authors:** Catherine R Knoverek, Upasana L Mallimadugula, Sukrit Singh, Enrico Rennella, Thomas E Frederick, Tairan Yuwen, Shreya Raavicharla, Lewis E Kay, Gregory R Bowman

## Abstract

Understanding the functional role of protein excited states has important implications in protein design and drug discovery. However, because these states are difficult to find and study, it is still unclear if excited states simply result from thermal fluctuations and generally detract from function or if these states can actually enhance protein function. To investigate this question, we consider excited states in β-lactamases and particularly a subset of states containing a cryptic pocket which forms under the Ω-loop. Given the known importance of the Ω-loop and the presence of this pocket in at least two homologs, we hypothesized that these excited states enhance enzyme activity. Using thiol labeling assays to probe Ω-loop pocket dynamics and kinetic assays to probe activity, we find that while this pocket is not completely conserved across β-lactamase homologs, those with the Ω-loop pocket have a higher activity against the substrate benzylpenicillin. We also find that this is true for TEM β-lactamase variants with greater open Ω-loop pocket populations. We further investigate the open population using a combination of NMR CEST experiments and molecular dynamics simulations. To test our understanding of the Ω-loop pocket’s functional role, we designed mutations to enhance/suppress pocket opening and observed that benzylpenicillin activity is proportional to the probability of pocket opening in our designed variants. The work described here suggests that excited states containing cryptic pockets can be advantageous for function and may be favored by natural selection, increasing the potential utility of such cryptic pockets as drug targets.

## Introduction

While it is well-established that proteins are dynamic molecules,[1] it is often unclear what these dynamics mean for function. An experimentally derived structural snapshot of a protein, such as a crystal structure, is frequently assumed to represent the (highest probability, lowest energy) ground state. This snapshot is also frequently assumed to be the functional state of the protein. In fact, rigidifying the active site, or increasing the probability of the ground state conformation, is often used as a design strategy for improving catalytic activity.[2, 3] In opposition to this common assumption, there are several compelling examples of functionally relevant excited states.[4–9] However, it is still unclear if excited states in general play a role in function.

Here, we consider an important class of excited states that contain a ‘cryptic’ pocket, or a pocket which is absent in the ligand-free, experimentally determined structure(s). These states are of particular interest because of the potential utility of cryptic pockets as drug targets.[10] These pockets provide a means to drug otherwise ‘undruggable’ proteins and a means to enhance a desired protein activity rather than just inhibit an undesired one.[11, 12] One concern, however, with the use of cryptic pockets as drug targets is that it is uncertain if there is a selective pressure to maintain the existence of a given pocket or if drug binding to that pocket could be trivially evolved away. This is at least partially because it is unknown if excited states containing cryptic pockets are simply a byproduct of the dynamic nature of proteins or if they play a bigger role in protein function.

Despite the many examples of systems which are known to contain cryptic pockets,[13–15] their functional relevance remains unclear, because these pockets are notoriously difficult to find and study. Identification of a cryptic pocket often requires simultaneous discovery of a ligand that binds to it.[16] Fortunately, recent advances in computational and experimental tools allow us to better identify and study these pockets.[1, 17] To increase sampling during molecular dynamics simulations, adaptive sampling methods like FAST[18] and replica exchange methods like SWISH[19] have been developed. To analyze these datasets, methods such as Markov state models (MSMs) [20] and exposons[21] have been developed. These computational tools can then be used to inform experimental methods like room temperature crystallography,[22] Nuclear Magnetic Resonance (NMR) relaxation techniques,[23–25] and thiol labeling assays.[26] Previous work using these methods has shown that many different kinds of proteins have cryptic pockets and that these pockets can be targeted with drugs to allosterically affect functional sites.[11, 12, 27, 28] However, it is still unclear if cryptic pockets have implications for function in the absence of ligand binding.

To explore the functional relevance of excited states containing cryptic pockets, we consider a set of class A β-lactamases. β-lactamases are enzymes that confer bacteria with antibiotic resistance by hydrolyzing β-lactam antibiotics, such as benzylpenicillin and cefotaxime. TEM β-lactamase, in particular, is an established model system for studying cryptic pockets. TEM has two known and well-characterized cryptic pockets. The first, which was found serendipitously during a drug screening campaign, is between helices 11 and 12.[16] The second, which was more recently identified in our laboratory,[21] forms when the Ω-loop undocks from the protein, so we call this pocket the Ω-loop pocket (Figure 1). The Ω-loop pocket was discovered in molecular dynamics simulations, confirmed using thiol labeling experiments, and subsequently shown to exert control over catalysis at the adjacent active site.[21] We know the Ω-loop structure is important as it is necessary for the deacylation of β-lactam antibiotics,[29] and we have previously shown that the total probability of conformations with a closed Ω-loop is predictive of cefotaxime activity.[30]

**Figure 1.**
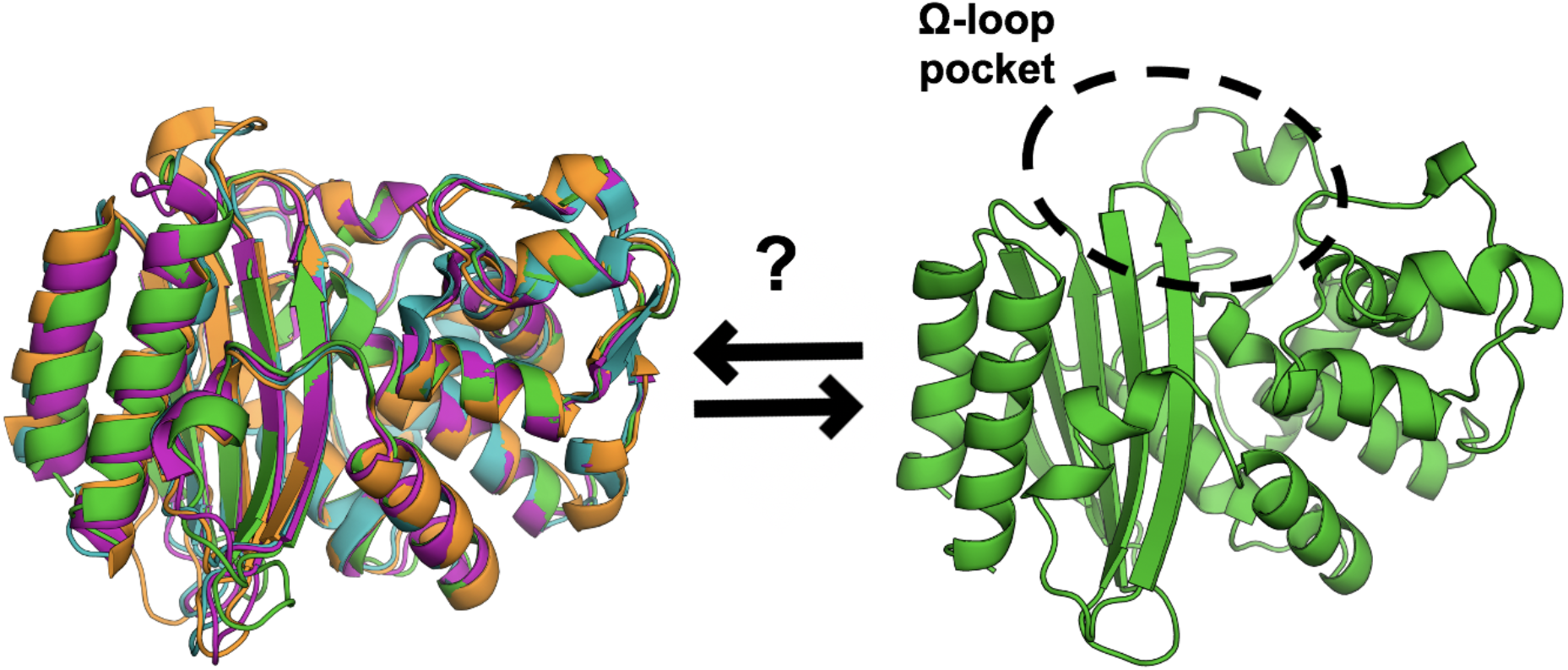
The Ω-loop pocket seen in TEM may open in other β-lactamase homologs. The structures of four β-lactamase homologs (left) overlay well. TEM (PDB: 1xpb) is shown in green, CTX-M-9 (PDB: 1ylj) is shown in cyan, MTB (PDB: 2gdn) is shown in orange, and GNCA (PDB: 4b88) is shown in magenta. The open Ω-loop pocket structure in TEM (right) was identified in molecular dynamics simulations.

As we have also found that the Ω-loop pocket is present in CTX-M-9 β-lactamase,[21] we hypothesize that this pocket may play a role in the enzyme’s function. To test this hypothesis, we first examine if the Ω-loop pocket is conserved across β-lactamase homologs and if the presence of the pocket is correlated with increased activity against classic β-lactam substrates. Here we mean conservation of the phenomenon of cryptic pocket opening, rather than conservation of the specific amino acid identities in that region of the protein. We then use activity data for TEM variants and combine NMR with molecular dynamics to gain insight into how the open Ω-loop pocket affects the hydrolysis reaction for different substrates. Finally, we design mutations to modulate the population of the open Ω-loop pocket to explicitly test whether pocket dynamics are predictive of enzymatic activity.

## Results/Discussion

### β-lactamase Homologs with Ω-Loop Pockets Hydrolyze Benzylpenicillin Faster

As a first step to determining if the Ω-loop cryptic pocket seen in TEM β-lactamase is functionally relevant, we examined if this pocket is conserved across β-lactamase homologs. Specifically, we examined MTB, the β-lactamase from *M. tuberculosis* encoded by the *blaC* gene, and GNCA, the predicted sequence for the last common ancestor of various Gram-negative bacteria as determined by ancestral sequence reconstruction.[31] Both of these proteins have the same topology as TEM (Figure 1, left), but only share about 50% sequence identity (Supplemental Figure 1). We know that the Ω-loop pocket is conserved in at least one homolog, CTX-M-9,[21] and wider conservation would suggest there is a selective pressure to maintain the pocket because it is playing a functional role. However, even if this pocket is not perfectly conserved, we hypothesize that the presence of the pocket may be correlated with enhanced enzyme activity. While closed Ω-loop conformations of TEM β-lactamase are predictive of increased cefotaxime activity,[30] there are often activity trade-offs in enzymes,[31] where increased activity for one substrate results in decreased activity for another. So, while a closed pocket may be beneficial for cefotaxime activity, an open pocket may be beneficial for a different substrate, such as benzylpenicillin.

To determine if the Ω-loop pocket is present in each homolog, we performed thiol labeling experiments that monitor the solvent exposure of a cysteine residue due to pocket opening. In these experiments, 5,5’-dithiobis-(2-nitrobenzoic acid), or DTNB, is added to the protein sample. DTNB covalently modifies exposed cysteine residues in a reaction that can be monitored as a change in absorbance at 412 nm over time. If a cysteine is buried inside of a pocket but then is exposed to solvent when the pocket opens, we observe an exponential increase in absorbance during the time scale of our experiment. We also ensure that the observed labeling rate is faster than the expected labeling due to protein unfolding, which is calculated from measuring the stability and/or unfolding rate. We have used this method previously to validate known,[26] and identify new,[21] cryptic pockets in TEM β-lactamase. As MTB and GNCA both have multiple native cysteine residues, with one cysteine located in the region of the Ω-loop pocket (Figure 2a-b), we added DTNB to each wild type homolog, monitored the change in absorbance, and used Beer’s law to calculate the number of cysteines labeling over time. When cysteine labeling was observed, we individually mutated out each native cysteine to determine which one is labeling in the wild type protein. For each homolog, we also measured the activity, beginning with the classic substrate, benzylpenicillin.

**Figure 2.**
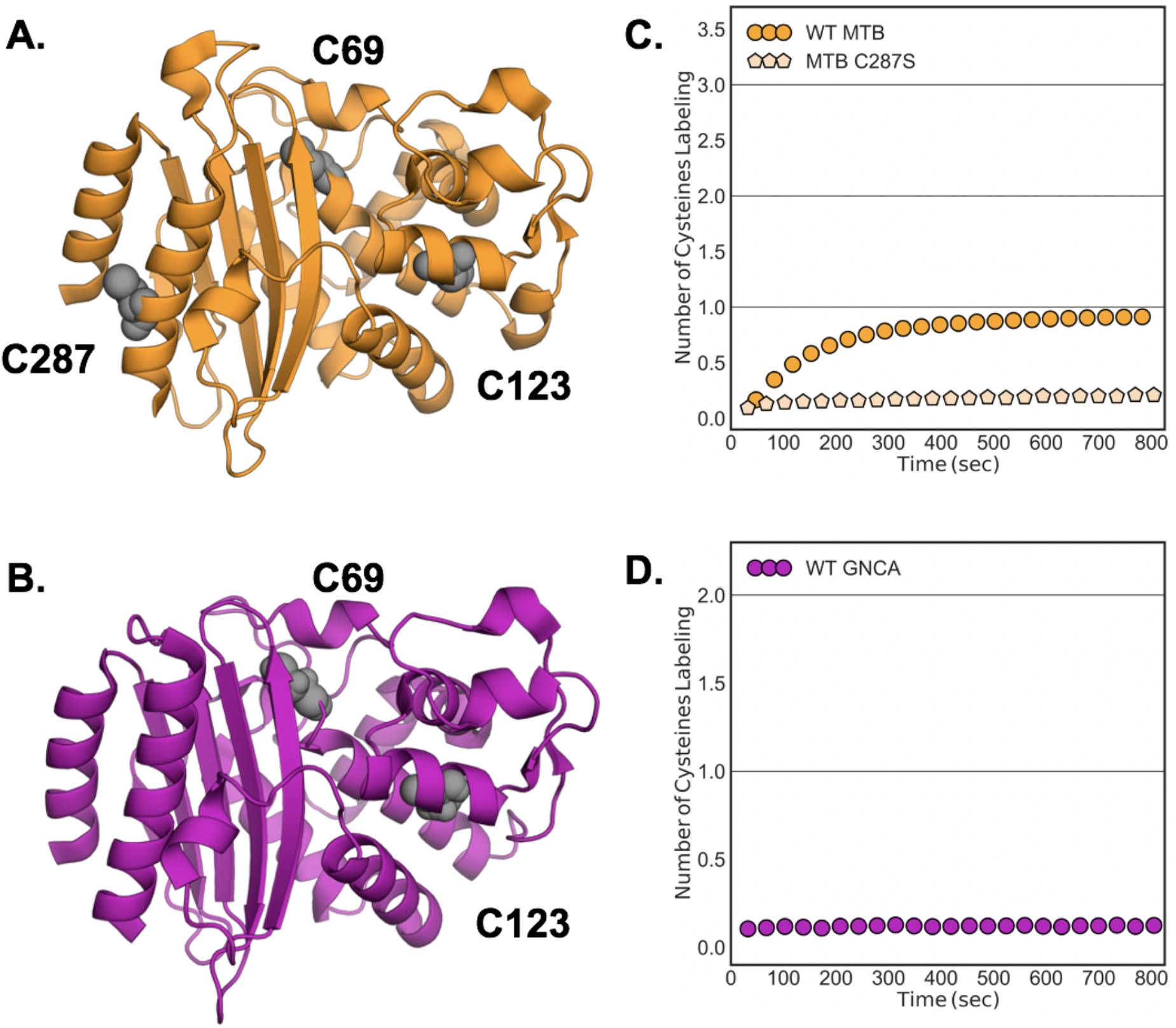
Labeling of MTB and GNCA β-lactamases suggest that neither protein open an Ω-loop pocket under the conditions tested. A-B. Structures of wild type (WT) MTB (PDB: 2gdn) and wild type GNCA (PDB: 4b88) are shown with native cysteine residues highlighted in gray. C69 is located in the region of the Ω-loop pocket for both proteins. C. The normalized DTNB labeling trace for WT MTB (orange circles) plateaus at one cysteine labeling. MTB C287S (light orange pentagons) shows significantly reduced labeling. D. The normalized labeling trace for WT GNCA shows no cysteine labeling.

For the conditions we tested, we find that neither MTB nor GNCA open a pocket in the region of the Ω-loop. Labeling of wild type MTB is well-fit by a single exponential (Supplemental Figure 2) and plateaus at the expected value for one cysteine labeling (Figure 2c). However, when we mutate out the cysteine residue in the Ω-loop pocket region, C69, the labeling overlays well with the wild type protein (Supplemental Figure 4). MTB C287S, however, displays significantly reduced labeling, suggesting that this is the cysteine that labels in the wild type protein. Thus, MTB does not open an Ω-loop pocket under the conditions tested here. Following the same procedure, we do not observe any significant labeling for wild type GNCA (Figure 2d). Again, the cysteine residue in the Ω-loop pocket region, C69, does not label, suggesting that GNCA also does not open an Ω-loop pocket under these conditions.

We also find, however, that the β-lactamase homologs with Ω-loop pockets display an increased ability to hydrolyze benzylpenicillin (Table 1). Penicillin binding proteins, which are unable to hydrolyze benzylpenicillin, lack the Ω-loop entirely.[29] The existence of this loop, and E166 in particular, in the MTB and GNCA homologs allows for the deacylation and completed hydrolysis of benzylpenicillin with moderate catalytic efficiency. However, TEM and CTX-M-9 both not only have the Ω-loop but are able to open a pocket in this region of the protein and have a corresponding increase in their ability to hydrolyze benzylpenicillin. If the substrate binding (~K_M_) but not the catalytic rate (k_cat_) was improved, that would suggest the pocket opening simply allows for the substrate to more easily enter the active site. However, both the overall catalytic efficiency and the catalytic rate are higher for TEM and CTX-M-9 than for MTB and GNCA. As a note, we do not expect the change in magnitude of the pocket open population to perfectly correspond to the change in catalytic rate. Our thiol labeling experiments monitor the equilibrium fluctuations of the apoenzyme while several β-lactamase conformations at various ligand binding stages are sampled during our activity assays. Also, because a more open pocket is beneficial, the increase in activity is not simply due to an increase in proximity of the catalytic residues. So, while the Ω-loop pocket is not conserved across all β-lactamase homologs, the pocket appears to be functionally relevant, because benzylpenicillin hydrolysis is increased when the pocket is present.

**Table 1.**
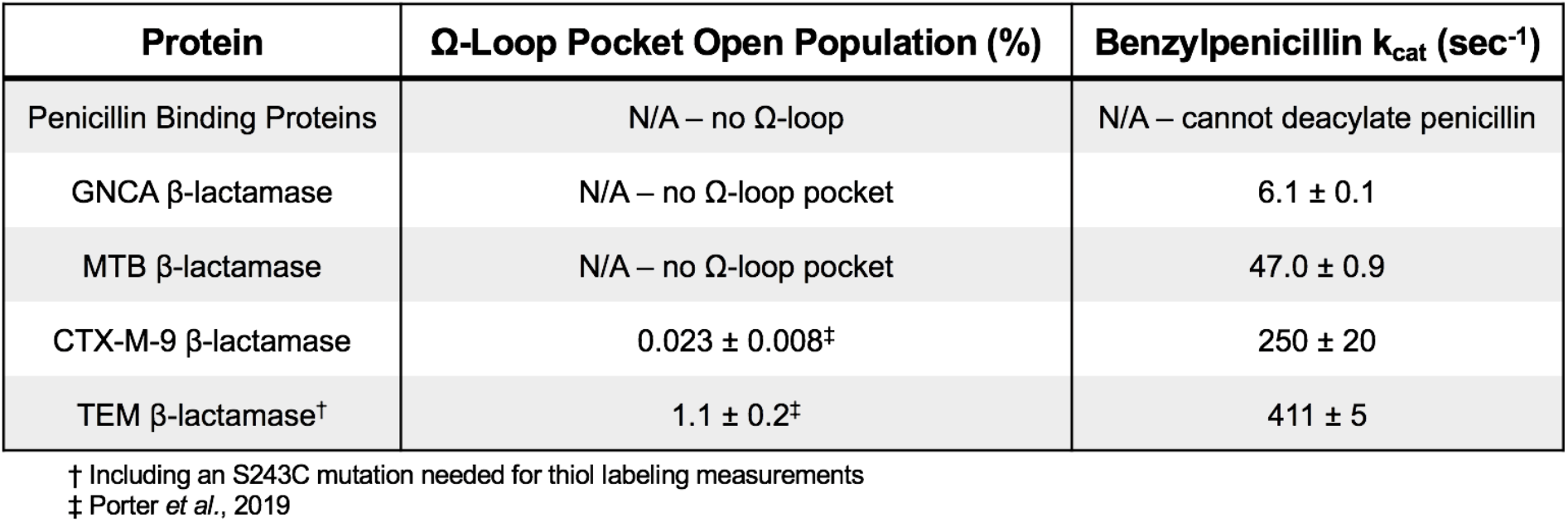
β-lactamase homologs with open Ω-loop pockets display increased catalytic rates against benzylpenicillin.

### TEM β-lactamase Variants with Higher Probabilities of Ω-Loop Pocket Opening Display Corresponding Increases in Benzylpenicillin Activity

To further test whether the Ω-loop pocket plays a role in the hydrolysis of benzylpenicillin, we focus our attention on a set of TEM variants. For TEM variants specifically, we have previously shown that there is a strong correlation between cefotaxime activity and the population of conformations with closed Ω-loop pockets.[30] So, we focus the rest of our study on TEM and hypothesize that an open Ω-loop pocket is detrimental for cefotaxime activity but beneficial for benzylpenicillin activity.

To begin investigating this hypothesis, we re-analyzed data from our previous study[30] where both benzylpenicillin and cefotaxime activity were measured for a number of TEM variants with mutations at clinically relevant positions. In the study, the benzylpenicillin activities were reported but not explored for their connection to the Ω-loop pocket dynamics. However, because of the correlation found between the Ω-loop closed conformations and cefotaxime activity, variants with decreased cefotaxime activity also have a higher population of conformations with an open Ω-loop pocket. Here, we examine how the catalytic rate for benzylpenicillin correlates to the catalytic efficiency against cefotaxime to determine if there is a trade-off between the two substrates, as this would suggest TEM variants with a higher population of open Ω-loop pocket conformations also have increased benzylpenicillin activity.

In fact, we do find a trade-off between benzylpenicillin and cefotaxime activity in the TEM variants examined here. We find that when TEM gains mutations that significantly increase cefotaxime activity, this corresponds to a decrease in catalytic rate for benzylpenicillin (Figure 3). As increased cefotaxime activity for these variants is correlated with a lower population of Ω-loop open conformations, the opposite is also true. Variants with a higher population of Ω-loop open conformations are correlated with increased benzylpenicillin activity. The variants form two distinct clusters, with the exception of TEM R164E/G238S and R164D/G238S (shown in gray), which display low activity against both substrates. Positions 164 and 238 are known to exhibit negative epitasis with one another,[32] so these variants likely have perturbed Ω-loop conformations that are deleterious for all substrates.

**Figure 3.**
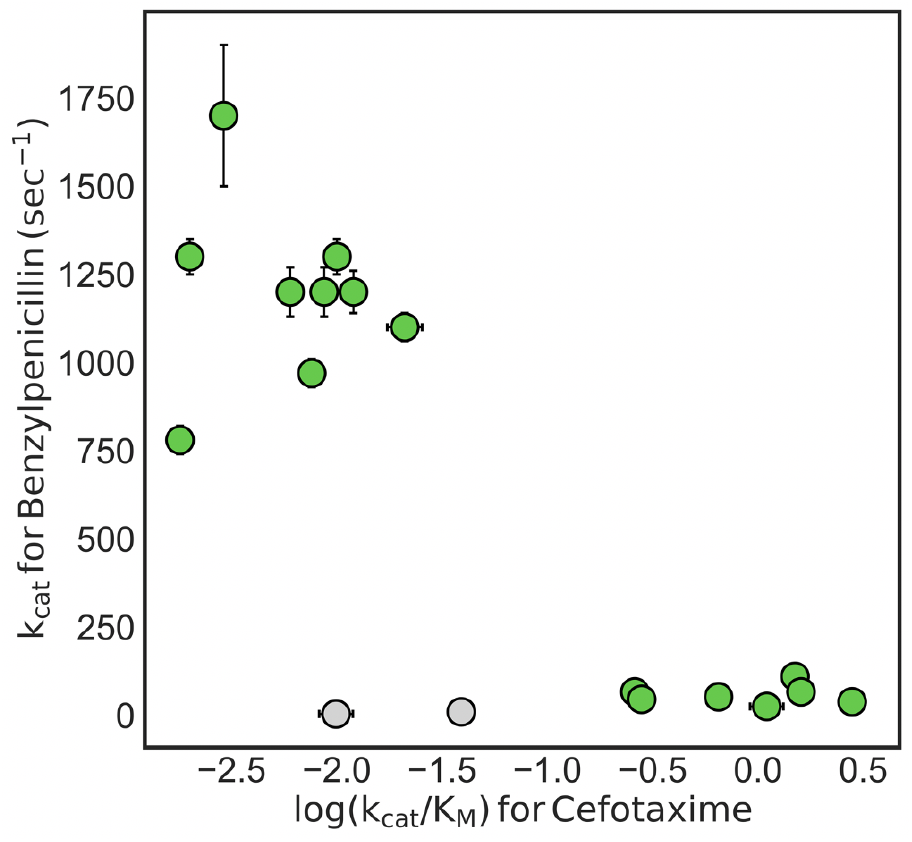
There is an inverse correlation between benzylpenicillin and cefotaxime activity for TEM variants. Each point represents a different TEM variant. Error bars represent standard error values from the fits. The variants shown in gray are R164E/G238S and R164D/G238S, which have decreased activity against both substrates. Data originally reported in [30].

### NMR and Simulations Provide Structural Insight into the TEM Ω-Loop Pocket Open Population Without the Need for a Mutation

To structurally characterize the open Ω-loop pocket population, we used a combination of NMR and molecular dynamics simulations. We performed an NMR relaxation technique called Chemical Exchange Saturation Transfer (CEST),[33, 34] which allows us to observe motions on the same timescale as our labeling experiments but without the need for a cysteine mutation. CEST can identify the presence of excited states and the rates of transitioning between states, thereby complementing our thiol labeling experiments. These NMR experiments also identify which residues contribute to the exchange between the ground state and excited state(s), which we used to inform the collective variable for metadynamics simulations. We then used conformations found during our metadynamics simulations to act as starting conformations for unbiased simulations on the Folding@home distributed computing platform.[28, 35] This “adaptive seeding” strategy allows us to access protein motions on the longer timescales of our labeling and NMR experiments while still preserving the thermodynamic and kinetic properties of the system. To analyze our simulation data, we used Correlation of All Rotameric and Dynamical States (CARDS)[36] followed by Principal Component Analysis (PCA) to identify the motions associated with the Ω-loop pocket and pull out exemplar structures for the pocket open and closed populations (Supplemental Figure 7).

Our CEST experiments provide further evidence for the TEM excited states predicted from our simulations and corroborate DTNB labeling data without the need for a mutation. Exchange was observed for TEM, suggesting the presence of an excited state population (Figure 4a). Many of the residues around the Ω-loop pocket report on to the exchange between the states, including L169, N170, G236, E240, G242, and S243 (Figure 4b, circles). The population of the excited state was determined to be 1.05 ± 0.03% (k_ex_ = 98 ± 6 sec^−1^), which is in excellent agreement with the open pocket population measured using thiol labeling (1.1± 0.2%). Furthermore, when we perform the same experiment on TEM with a mutation that abolishes the Ω-loop pocket (R241P, see data below), the dynamics monitored by CEST disappear (Figure 4b, pentagons) as well as faster timescale dynamics monitored by relaxation dispersion (Supplemental Figure 6). These data suggest that CEST reports on the Ω-loop pocket open population. We then used the residues identified in these experiments as being dynamic on the millisecond to second timescale to define the collective variable for metadynamics simulations, which in turn identified seed conformations for unbiased simulations.

**Figure 4.**
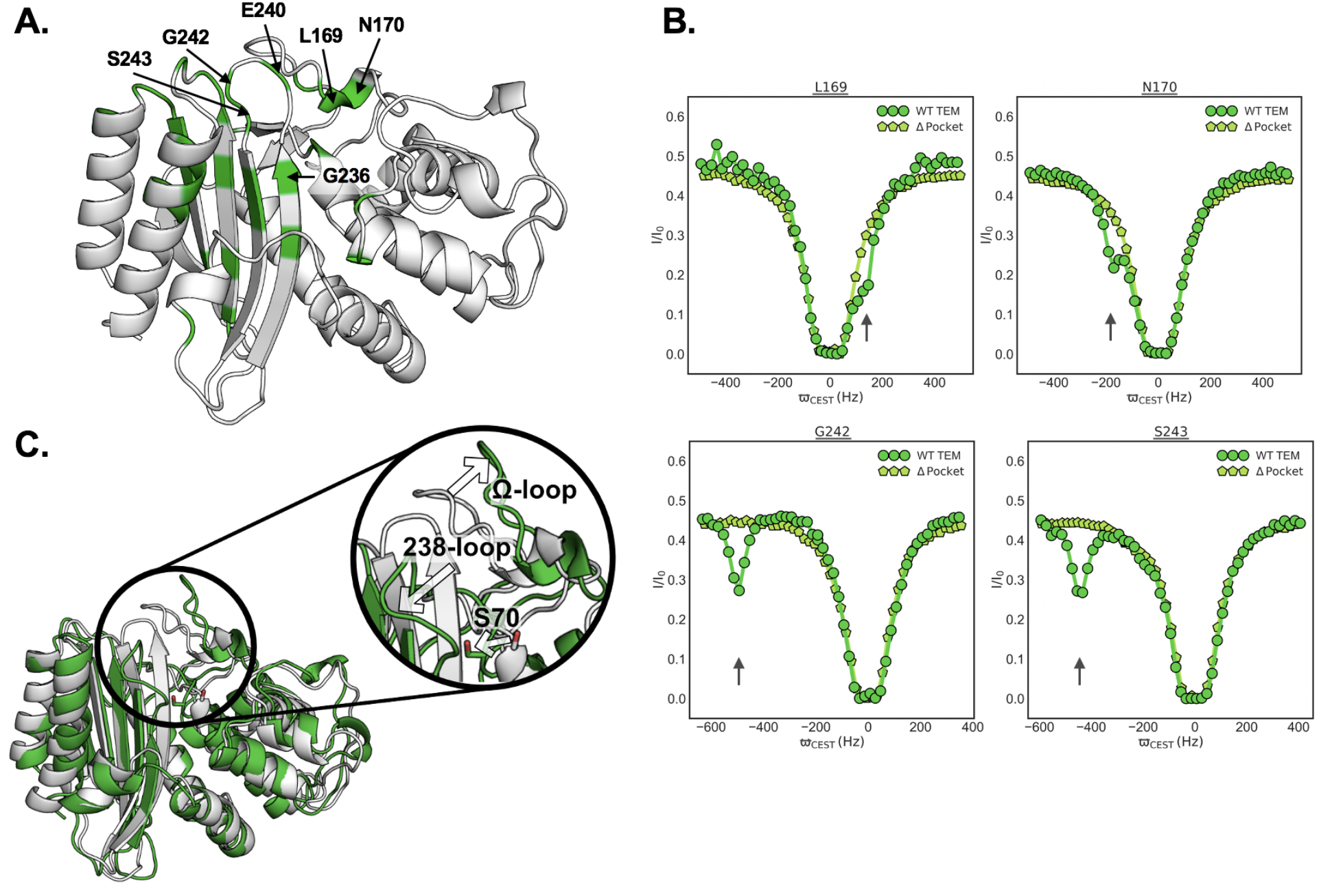
Structural insight into the TEM open Ω-loop pocket population identifies conformational changes in the 238-loop and catalytic S70. A. Highlighted in green on the wild type (WT) TEM structure (PDB: 1xpb) are the residues showing conformational exchange as established by minor dips in the ^15^N CEST profiles. Residues around the Ω-loop pocket are annotated. B. ^15^N CEST profiles for wild type TEM (green circles) and TEM R241P, a variant with no Ω-loop pocket, (light green pentagons) are shown for residues in the Ω-loop pocket. Minor dips seen in the wild-type protein are highlighted with arrows. These experiments informed molecular dynamics simulations that identified the structures shown in C. C. The closed pocket conformation (white) is similar to the crystal structure. The open pocket conformation (green) has an open Ω-loop, an open 238-loop, and a buried catalytic S70 (shown in sticks).

Analysis of our simulations show an Ω-loop pocket open state with an open Ω-loop and open 238-loop (Figure 4c), in good agreement with the NMR experiments reported here as well as our previous thiol labeling of this pocket.[21] We find that the Ω-loop closed conformation looks very similar to the crystal structure, as expected. We also find that Ω-loop pocket opening is correlated with burial of the catalytic serine (S70), making the residue no longer available to bind substrate (Supplemental Figure 8). For TEM β-lactamase, which hydrolyzes benzylpenicillin with an efficiency approaching the diffusion limit,[37] a buried S70 following deacylation may prevent re-acylation, and an open Ω-loop pocket may promote product release by reducing protein-product contacts. On the other hand, wild type TEM poorly degrades cefotaxime due to a high K_M_ and low acylation rate. In this case, a more closed Ω-loop pocket would increase protein-substrate contacts, and an available S70 would be beneficial for substrate binding and subsequent acylation.

### Variants Designed to Modulate the Dynamics of the Ω-Loop Pocket Predictably Affect Function

Finally, we explicitly test our model by designing variants to either close or open the Ω-loop pocket, assessing the impact on pocket opening via thiol labeling experiments, and observing the effects on both benzylpenicillin and cefotaxime hydrolysis functions. We aimed to make mutations in TEM that are not known to be involved in catalysis directly but that would affect the Ω-loop pocket dynamics. Thus, changes in the pocket dynamics resulting in predictable changes in activity would support our understanding of how the TEM Ω-loop pocket is connected to function.

Towards that end, we selected Ω-loop pocket mutations based off of a previous study[38] that measured the fitness effects of every single point mutation in TEM in the presence of either ampicillin or cefotaxime. An E240D mutation produced a positive fitness effect in the presence of ampicillin. While a mutation to lysine at this position is known to be clinically important, the charge conserving mutation to aspartic acid at this position has not been seen clinically. We rationalized that mutating this position to an aspartic acid would destabilize closed Ω-loop conformations due to its shorter hydrocarbon chain leading to lower hydrophobicity and reduced ability to screen its charged, acid group by positioning it into the solvent. Following this logic, we hypothesized that the E240D mutation opens the Ω-loop pocket in TEM, which in turn increases benzylpenicillin activity. On the other hand, a R241P mutation produced a large positive fitness effect in the presence of cefotaxime. This position is not known to be clinically important, and we reasoned that removal of a charged amino acid might stabilize closed conformations and introduction of a proline, which has fewer available dihedral angles, would destabilize open conformations. Thus, we hypothesized that the R241P mutation closes the Ω-loop pocket in TEM, which increases cefotaxime activity and decreases benzylpenicillin activity.

As predicted, we find that the E240D mutation opens the Ω-loop pocket, decreases cefotaxime activity, and increases benzylpenicillin activity (Figure 5). The thiol labeling rates for TEM E240D are faster than wild type, indicating a more open Ω-loop pocket (population = 4 ± 1%, EXX regime). As expected, the catalytic efficiency of E240D for benzylpenicillin increases. The relatively small increase is also expected as TEM is very close to the diffusion limit for this substrate. We also see the corresponding decrease in cefotaxime activity for this mutant.

**Figure 5.**
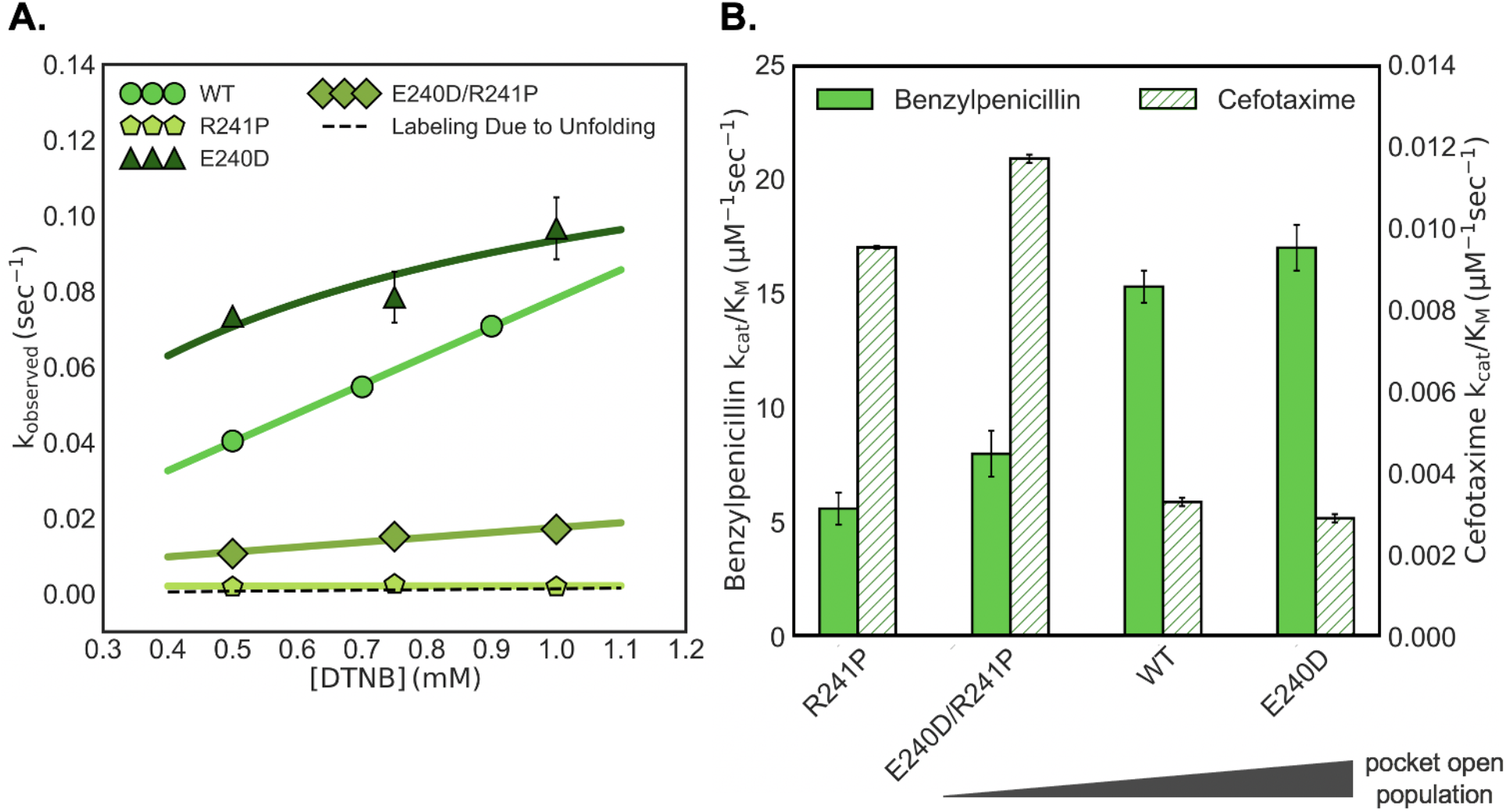
Mutations in TEM designed to alter the open Ω-loop pocket population lead to predictable changes in benzylpenicillin and cefotaxime activity. A. The observed labeling rate as a function of DTNB concentration is shown for wild type (WT) TEM (circles), TEM E240D (triangles), TEM R241P (pentagons), and TEM E240D/R241P (diamonds). Higher labeling rates are due to a higher open Ω-loop pocket population. The dashed line represents the expected labeling for wild type TEM due to the unfolded population. Error bars represent the standard deviation of three measurements. B. Benzylpenicillin (solid) and cefotaxime (striped) activity is shown for each variant. Error bars are the result of bootstrapping analysis.

We find that the R241P mutation also behaves as expected. The thiol labeling rates for TEM R241P are much slower than wild type. In fact, they are on the order of magnitude as those expected for labeling due to protein unfolding, thus suggesting that R241P abolishes the Ω-loop pocket in TEM. As a result, the activity of R241P against benzylpenicillin decreases, and we find a corresponding increase in the catalytic efficiency against cefotaxime, which is consistent with our model and suggests that the mutation does not just simply break the enzyme.

When we introduce the E240D mutation into the background of R241P, we find that the opening of the Ω-loop pocket is rescued (Figure 5a, population = 0.19 ± 0.01%, EX2 regime). The corresponding benzylpenicillin activity for TEM E240D/R241P is, as expected, higher than that of TEM R241P. It is also lower than that of wild type TEM, consistent with the stronger effect of the R241P mutation than that of the E240D mutation in the wild-type background. These data support our hypothesis that a higher open Ω-loop pocket population in TEM is correlated with higher benzylpenicillin activity. TEM E240D/R241P also has increased cefotaxime activity compared to wild type TEM, which is consistent with our model that closed Ω-loop conformations are beneficial for cefotaxime activity. However, the cefotaxime activity of TEM E240D/R241P, which has a low Ω-loop pocket open population, is greater than the cefotaxime activity for TEM R241P, which does not have an Ω-loop pocket. This suggests that, even though closed Ω-loop pocket conformations are beneficial for cefotaxime activity, a low open pocket population may be more advantageous than no pocket at all. Taken together, the results presented here suggest that the Ω-loop pocket in β-lactamase plays a role in function, as excited states containing an open pocket improve activity.

## Conclusions

We hypothesized that β-lactamase excited states play a role in function by increasing enzyme activity. Specifically, we investigated whether excited states containing the Ω-loop cryptic pocket enhance benzylpenicillin activity. We found that the Ω-loop pocket seen in TEM and CTX-M-9 β-lactamase is not conserved in the MTB and GNCA homologs, but that homologs with the pocket have higher catalytic rates for the hydrolysis of benzylpenicillin. Focusing our study on TEM β-lactamase, we found that variants with larger open Ω-loop pocket populations also have higher benzylpenicillin catalytic rates. To gain structural insight into the TEM open Ω-loop pocket population, we performed NMR CEST experiments and NMR-guided molecular dynamics simulations. Lastly, we designed mutations to modulate the dynamics of the Ω-loop pocket in TEM and observed that the probability of pocket opening is predictive of benzylpenicillin activity. Our results demonstrate our understanding of how excited states containing the Ω-loop cryptic pocket are connected to β-lactamase function and provide further evidence for the hypothesis that functionally relevant conformations are sampled during equilibrium fluctuations (*i.e*., in the apoenzyme). This work also demonstrates that cryptic pocket dynamics can be modulated with mutations, setting up future studies to elucidate the sequence determinants of these pockets, and suggests that cryptic pockets may be under positive selective pressure, increasing their potential utility as drug targets.

## Author Contributions

CRK, LEK, and GRB designed the research. CRK, ULM, SS, ER, TF, TY, and SR performed research. CRK, ULM, SS, ER, and TF analyzed the data. CRK and GRB wrote the paper.

## Acknowledgements

We would like to thank the community scientists of Folding@home for donating their computing resources. This work was funded by National Institutes of Health grant R01GM12400701 (GRB), National Science Foundation CAREER Award MCB-1552471 (GRB), Canadian Institutes of Health Research FDN-503573 (LEK), and the Natural Sciences and Engineering Research Council of Canada (LEK). GRB holds a Career Award at the Scientific Interface from the Burroughs Wellcome Fund and a Packard Fellowship for Science and Engineering from the David and Lucile Packard Foundation. LEK holds a Canada Research Chair in Biochemistry.

## Methods

### Mutagenesis and protein purification

We previously cloned the genes for TEM and CTX-M-9 β-lactamase into pET24-b plasmids (Life Technologies) for inducible protein expression under the T7 promoter.[21, 30] Both plasmids use kanamycin resistance for selection and the TEM plasmid contains the OmpA signal sequence for periplasmic export. MTB and GNCA were cloned into pET28 plasmids by Genewiz for inducible protein expression under the T7 promoter. Both plasmids use kanamycin resistance for selection. The MTB plasmid contains an N-terminal 6x His tag with thrombin cleavage site, and the GNCA plasmid contains the OmpA signal sequence for periplasmic export. We created protein variants using site-directed mutagenesis and verified the mutations via DNA sequencing. For expression, we transformed our desired plasmid into BL21(DE3) cells (Intact Genomics) and grew cultures to an OD_600_ of 0.6 before we induced protein expression by adding 1 mM IPTG. TEM, MTB, and GNCA were expressed overnight at 18°C, while CTX-M-9 was expressed for at least three hours at 37°C.

We purified TEM and GNCA using our previously described protocol[30] that isolates the protein from the periplasm using the following osmotic shock lysis protocol. We harvested cells and resuspended them in 30 mM Tris, pH 8.0 with 20% sucrose. After centrifugation, we resuspended the cells in 5 mM MgSO_4_ at 4°C. After another centrifugation, we dialyzed the supernatant against 20 mM sodium acetate, pH 5.5 overnight at 4°C. We centrifuged the dialysis contents to remove any insoluble protein and then purified using cation exchange chromatography (BioRad UNOsphere Rapid S column) with an NaCl gradient. The final purification step was size exclusion chromatography (BioRad ENrich SEC 70 column), and we stored the purified protein at 4°C in 20 mM Tris, pH 8.0.

We purified CTX-M-9 as previously described[21] by isolating the protein from inclusion bodies using the following protocol. We harvested cells and resuspended them in 20 mM sodium acetate, pH 5.5 and froze them at −80°C at least overnight. We then thawed the cells and lysed them via sonication. We centrifuged the lysate, and resuspended the pellet in 20 mM sodium acetate, pH 5.5 + 9 M urea overnight. After centrifugation, we refolded the protein by adding it drop-wise to buffer with no urea while gently stirring. We removed aggregated protein by centrifugation and dialyzed the supernatant against 20 mM sodium acetate, pH 5.5 overnight at 4°C. After centrifuging the dialysis contents to remove any insoluble protein, we then purified using cation exchange chromatography (BioRad UNOsphere Rapid S column) with an NaCl gradient. The final purification step was size exclusion chromatography (BioRad ENrich SEC 70 column), and we stored the purified protein at 4°C in 20 mM Tris, pH 8.0.

We purified MTB by isolating the protein from the cytoplasm using the 6x His tag and the following protocol. Cells were harvested and resuspended in 25 mM Tris, pH 7.5 + 300 mM NaCl and frozen at −80°C overnight. Cells were then thawed and lysed via sonication. The lysate was centrifuged to remove cell debris and the supernatant was loaded onto a Ni-NTA agarose column. Elution peak fractions were then dialyzed against 25 mM Tris, pH 7.5 + 300 mM NaCl overnight at 4°C. The dialysis contents were centrifuged to remove any insoluble protein before the 6x His tag was removed by thrombin cleavage. The reaction was carried out overnight at room temperature while stirring. The reaction contents were centrifuged to remove any insoluble protein and then cleaved protein was isolated by collecting the flow-through of a Ni-NTA agarose column run. The final purification step was size exclusion chromatography (BioRad ENrich SEC 70 column), and the purified protein was stored at 4°C in 20 mM Tris, pH 8.0.

### Labeling assays

For pocket determination, we labeled 5-30 μM protein with 2 mM DTNB (Ellman’s reagent, Thermo Scientific) in 20 mM Tris, pH 8.0 at 25°C until completion. We monitored the reaction via a change in absorbance at 412 nm over time using a Cary 100 UV–Vis spectrophotometer (Agilent Technologies). Each measurement was performed in triplicate. We determined the number of cysteines that labeled by first using a series of exponentials fit to the data and then by normalizing the signal using the known protein concentration and Beer’s Law (below). Here, *l* is the pathlength of the cuvette, which is one cm, and *ε* is the extinction coefficient of TNB at 412 nm, which is 14,150 M^−1^ cm^−1^.[39]

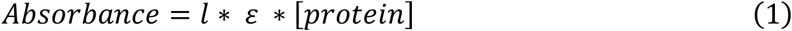

For labeling of the Ω-loop pocket of the TEM variants, we labeled 10 μM protein with various concentrations of DTNB (Ellman’s reagent, Thermo Scientific) in 20 mM Tris, pH 8.0 at 25°C until completion. We introduced an S243C mutation in order to observe pocket opening via labeling, as previous described.[21] We monitored the reaction via a change in absorbance at 412 nm over time using a Cary 100 UV–Vis spectrophotometer (Agilent Technologies) and performed measurements at each DTNB concentration in triplicate. Next, we fit a single exponential equation to the data to obtain the observed rate constants and plotted these values as a function of DTNB concentration. We fit the Linderstrøm-Lang model[40] (below) to the observed rate constants as a function of DTNB concentration to obtain the open pocket population, and obtained error using bootstrapping. We find that the TEM variants reported in this study displayed labeling in the EXX (full expression) or EX2 regime. The EX2 regime is the limiting case when the rate of pocket closing is much faster than the intrinsic rate of labeling, and the observed rate depends linearly on the DTNB concentration.

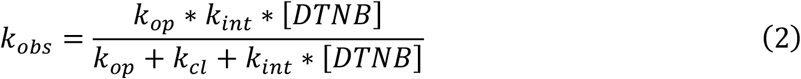

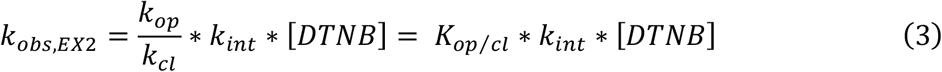

Here, *k_obs_* is the observed labeling rate, *k_op_* is the rate of pocket opening, *k_cl_* is the rate of pocket closing, and *k_int_* is the intrinsic labeling rate. We previously measured the intrinsic labeling rate (6.83 sec^−1^ mM^−1^) by performing the same labeling experiment but on a five amino acid peptide of protein sequence containing the cysteine of interest.[21] The equilibrium constant for pocket opening (*K_op/cl_*) should be greater than the equilibrium constant for unfolding, which was measured using urea denaturation experiments (see supplemental methods). We used the equilibrium constant for wild type TEM unfolding in equation (3) to calculate *k_obs_* as a function of DTNB concentration due to unfolding (dashed line in main text Figure 5a).

### Activity assays

We measured the initial velocity (*v_i_*) of antibiotic degradation by β-lactamase at 25°C via a change in absorbance (232 nm for benzylpenicillin, 262 nm for cefotaxime) using a Cary 100 UV–Vis spectrophotometer (Agilent Technologies). The substrate (5-200 μM) was incubated at 25°C for 5 min before addition of the protein. We diluted purified protein to a final concentration no greater than 200 nM. Our activity buffer was 50 mM potassium phosphate, pH 7.0 with 10% glycerol, and we measured each substrate concentration in triplicate. For benzylpenicillin, the Michaelis-Menton equation (below) was fit to the initial velocity as a function of the substrate concentration to determine individual catalytic rate (*k_cat_*) and Michaelis constant (*K_M_*) values. Here, [*E*] is the total enzyme concentration, and [*S*] is the total substrate concentration. For cefotaxime, the K_M_ was too high to reach maximum velocity, so a line (below) with a slope equal to 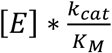 was fit to the data. Error for the fit parameters was determined using bootstrapping.

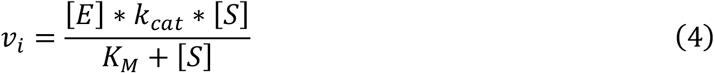

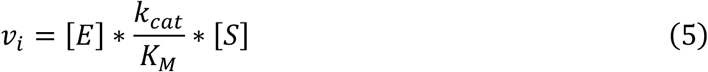

### NMR CEST experiments

We recorded all experiments on a Bruker AVANCE III HD 18.8 T spectrometer equipped with a cryogenically cooled, x,y,z pulsed-field gradient triple-resonance probe. We recorded each D-CEST[41, 42] experiment (30°C) as a pseudo-3D matrix, where each 2D spectrum were obtained as a function of the position of weak B_1_ perturbations applied at discrete frequencies overt the chemical shift range of the probed nucleus. We applied a DANTE excitation scheme,[43] which perturbs multiple regularly spaced frequencies at the same time, thereby decreasing the frequency range that must be explored over regular CEST approaches. In all cases, we calibrated the strength of the B_1_ field using a nutation experiment, as described previously.[44]

We acquired ^15^N D-CEST data as previously described[41] using 1 s DANTE excitation trains of square pulses (~7° flip angle, 2.5 kHz B_1_ field) and an interpulse delay of 2, 1, and 0.667 ms, resulting in effective B_1_ fields of about 10, 20, and 30 Hz. We sampled CEST profiles in 51 steps, with increments of 10, 20, and 30 Hz, extending over frequency ranges of 500 (2 ms interpulse delay), 1000 (1ms), and 1500 Hz (0.667 ms), respectively. We extracted the position of the minor dips, exchange rate, and population of the excited state by fitting a two-state model of chemical exchange to the CEST data as described in detail previously.[32] Errors were estimated using the bootstrapping.

### Molecular dynamics simulations and analysis

We prepared systems as previously described,[20] using GROMACS software[45] and the Amber03 force field.[46] We solvated in TIP3P water[47] and energy minimized using the steepest descent algorithm. We used the V-rescale thermostat to maintain a fixed temperature of 300K and the Berendsen barostat to bring the pressure up to one bar. Mutations were introduced into the starting structure using PyMol.

We ran 200 nanoseconds of metadynamics simulations on each variant using the PLUMED plugin on GROMACS[48] and defined our collective variable (*s*) using the backbone torsional angles of the 238-loop. This collective variable is expressed using the equation below.

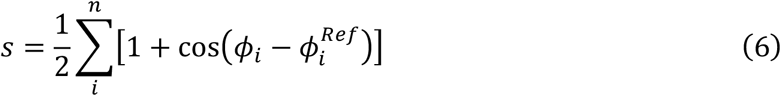

Here, *ϕ_i_* is the torsion angle of dihedral *i* of the current frame in the simulation and 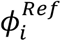 is a reference torsion angle defined by the corresponding dihedral in the crystal structure. This summation is done across *n* dihedrals. We computed our collective variable using the *ϕ* and *ψ* angles of residues that underwent CEST, namely residues 237 through 243. Gaussians were added every 2 picoseconds with a height of 1.0 kJ/mol and a width of 0.05.

We clustered our metadynamics simulations using a hybrid k-centers/k-medoids algorithm[49] to generate 220 representative seed conformations, using a cluster radius cutoff of 1.2 Å. We then used these conformations to collect a total of 100.7 microseconds of unbiased simulations on our Folding@home distributed computing platform.[50]

To analyze our simulation data, we first built a Markov state model (MSM) using our Enspara software.[51] We again clustered using a hybrid k-centers/k-medoids algorithm and used a cluster radius cutoff of 1.2 Å, which resulted in 9877 states. A pseudo-count was added to each element in the transition counts matrix to prevent sampling artifacts from influencing the transition probabilities.

We then used the CARDS methodology[36] to compute the holistic communication (*I_H_*(*X,Y*)) for every pair of dihedrals *X* and *Y* using the equation below.

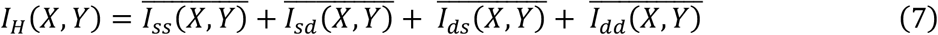

Here, 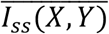 is the normalized mutual information between the structure (i.e., rotameric state) of dihedral *X* and the structure of dihedral *Y*, 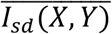 is the normalized mutual information between the structure of dihedral *X* and the dynamical state of dihedral *Y*, 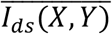 is the normalized mutual information between the dynamical state of dihedral *X* and the structure of dihedral *Y*, and 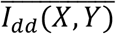 is the normalized mutual information between the dynamical state of dihedral *X* and the dynamical state of dihedral *Y*. The mutual information (*I*) is described by the equation below.

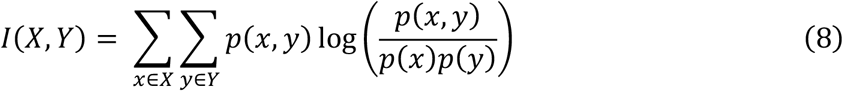

Here, *x* ∈ *X* refers to the set of possible states that dihedral *X* can adopt, *p*(*x*) is the probability that dihedral *X* adopts state *x*, and *p*(*x,y*) is the joint probability that dihedral *X* adopts state *x* and dihedral *Y* adopts state *y*. We computed normalized mutual information using the maximum possible mutual information, known as the channel capacity, for any specific mode of communication. We then computed a community network using affinity propagation,[52] with a damping parameter of 0.8. We generated the final allosteric network by filtering the community network using the Marginal Likelihood Filter (MLF)[53] to capture the top 5% of edges.

Finally, we applied principal component analysis (PCA) to the distances between the Cβ atoms of every pair of residues in the community containing the Ω-loop and catalytic S70. We projected our MSM onto principal components 1 and 3 (PC1 and PC3) and pulled out exemplar structures by estimating the population-weighted centroid of the two minima.

## Supplemental Information

### Supplemental Figures

**Supplemental Figure 1.**
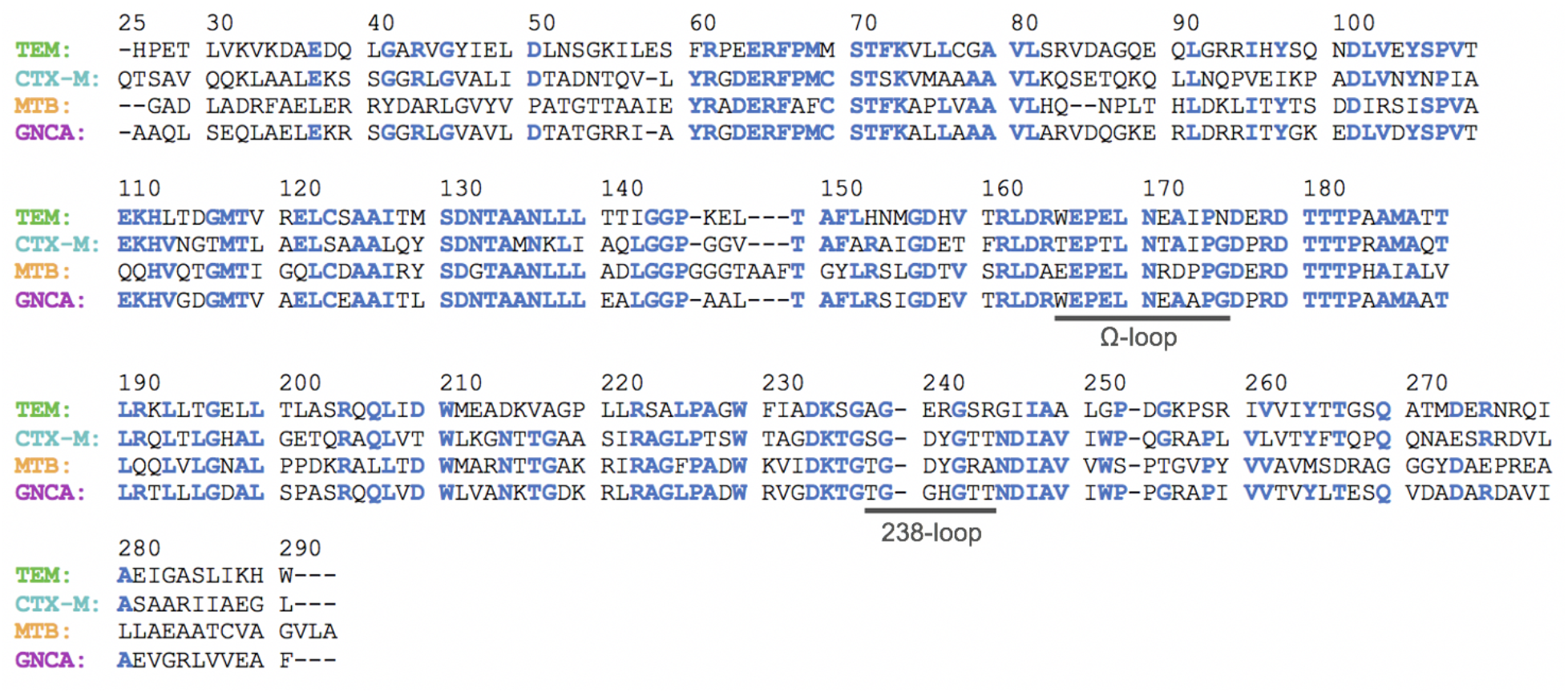
β-lactamase homologs have the same topology, but only share about 50% sequence identity. The sequence alignment for the four homologs discussed in this study is shown here, with conserved residues shown in blue. The Ω-loop and 238-loop sequences are underlined.

**Supplemental Figure 2.**
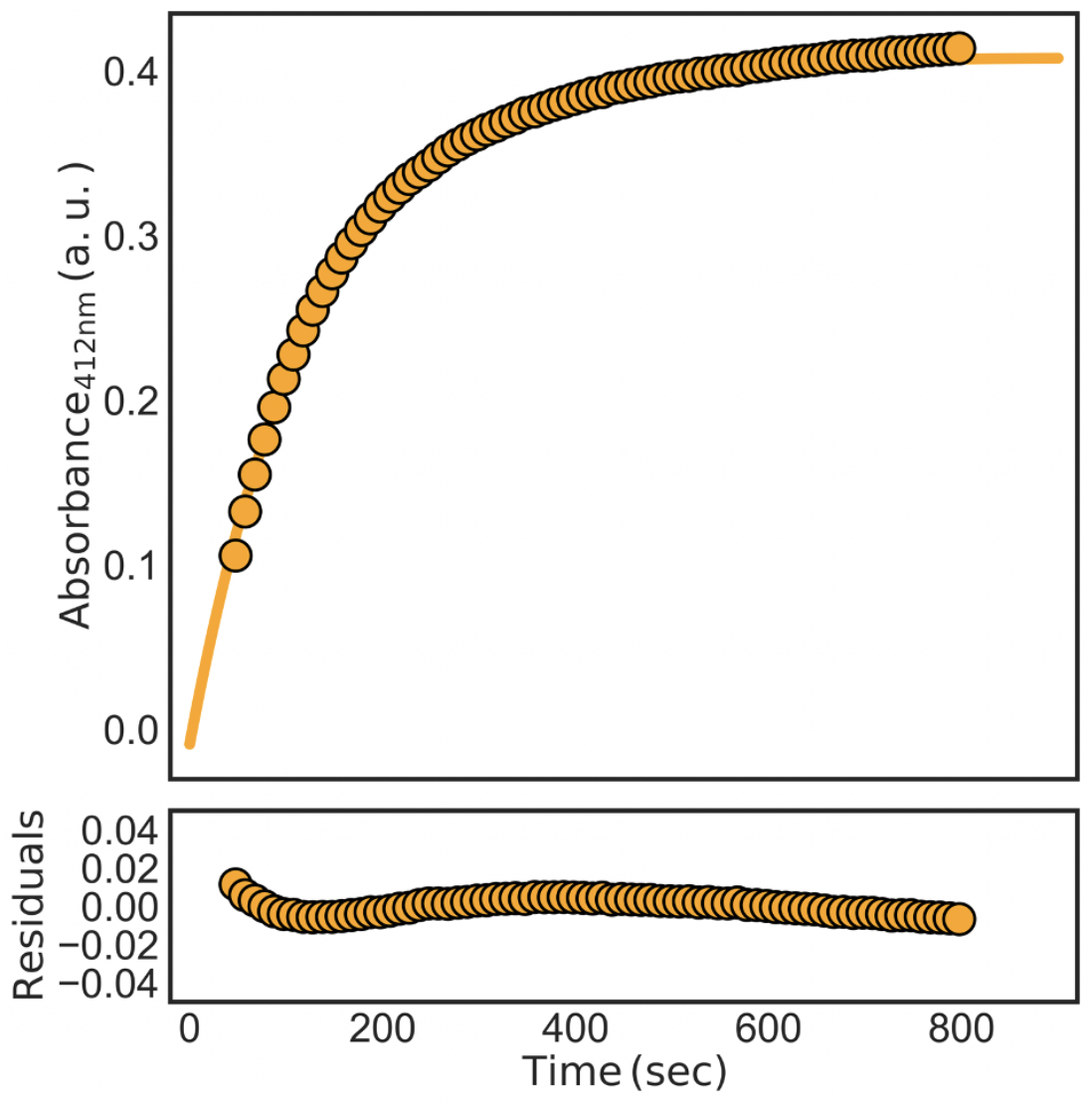
Labeling of WT MTB is well-fit by a single exponential. Shown here is the average labeling trace for 30 μM protein and 2 mM DTNB. Raw data is shown as circles and the fit is shown as a solid line. Below are the residuals for the fit.

**Supplemental Figure 3.**
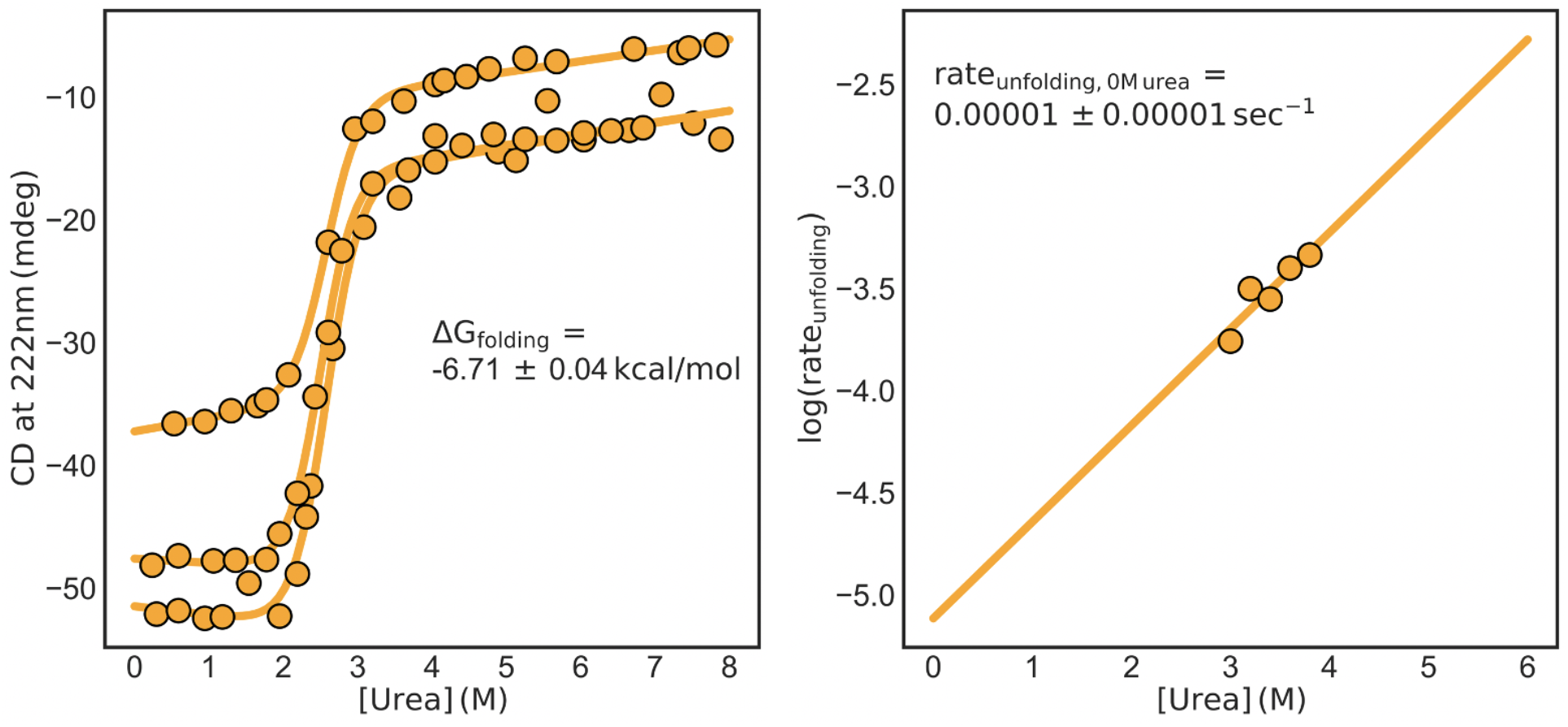
Labeling of WT MTB was not due to protein unfolding. (Left) Urea denaturation was followed by circular dichroism. Error is reported as the standard deviation of three replicate experiments. (Right) The unfolding rate was monitored by circular dichroism and then plotted as a function of urea concentration. The y-intercept of the fit line represents the unfolding rate in the absence of urea.

**Supplemental Figure 4.**
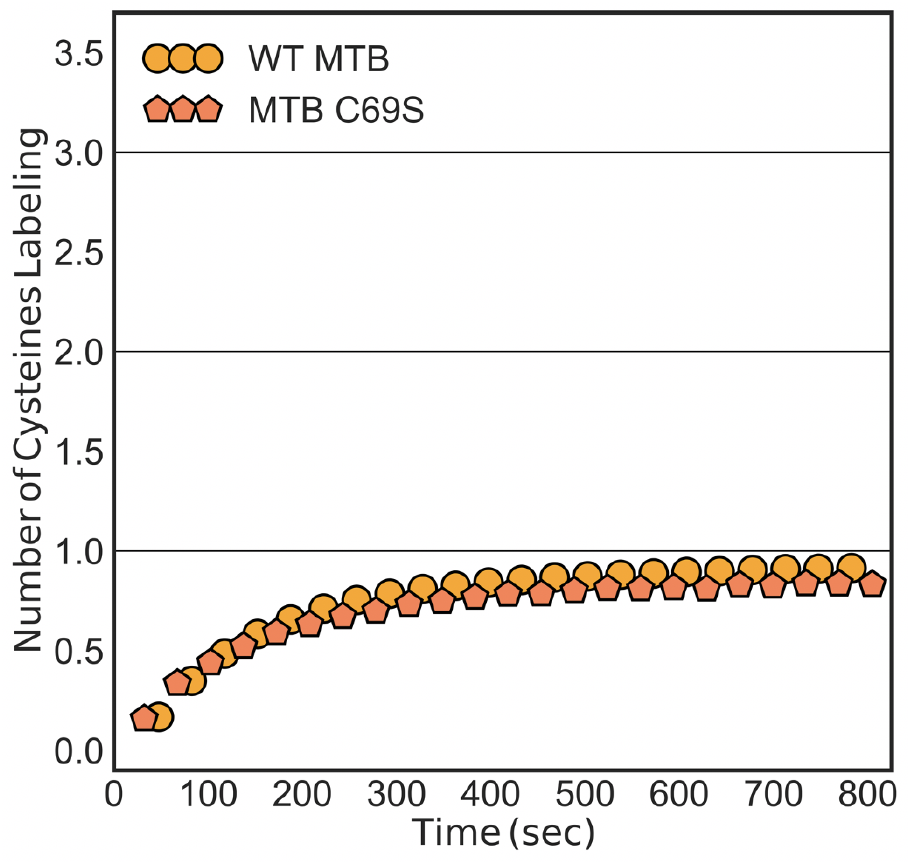
Mutating out the cysteine residue in the Ω-loop pocket region, C69, does not reduce labeling. The normalized DTNB labeling of MTB C69S (coral pentagons) overlays well with the labeling of WT MTB (orange circles), both which plateau at one cysteine labeling.

**Supplemental Figure 5.**
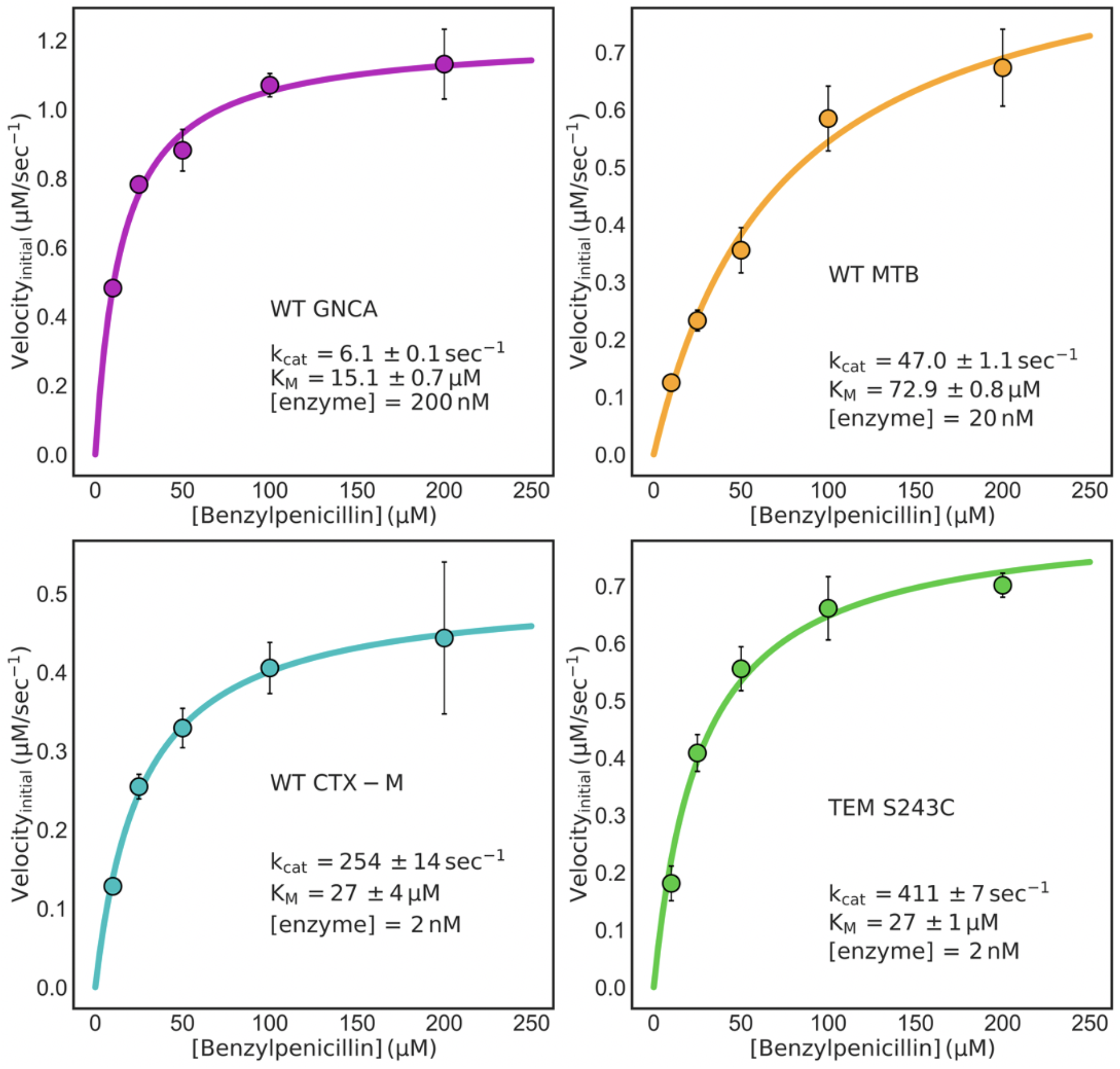
β-lactamases with Ω-loop pockets have higher catalytic rates against benzylpenicillin. The full Michaelis-Menten equation was fit to the data. Error bars are shown as the standard deviation of three replicate measurements. Error for each parameter was determined using bootstrapping.

**Supplemental Figure 6.**
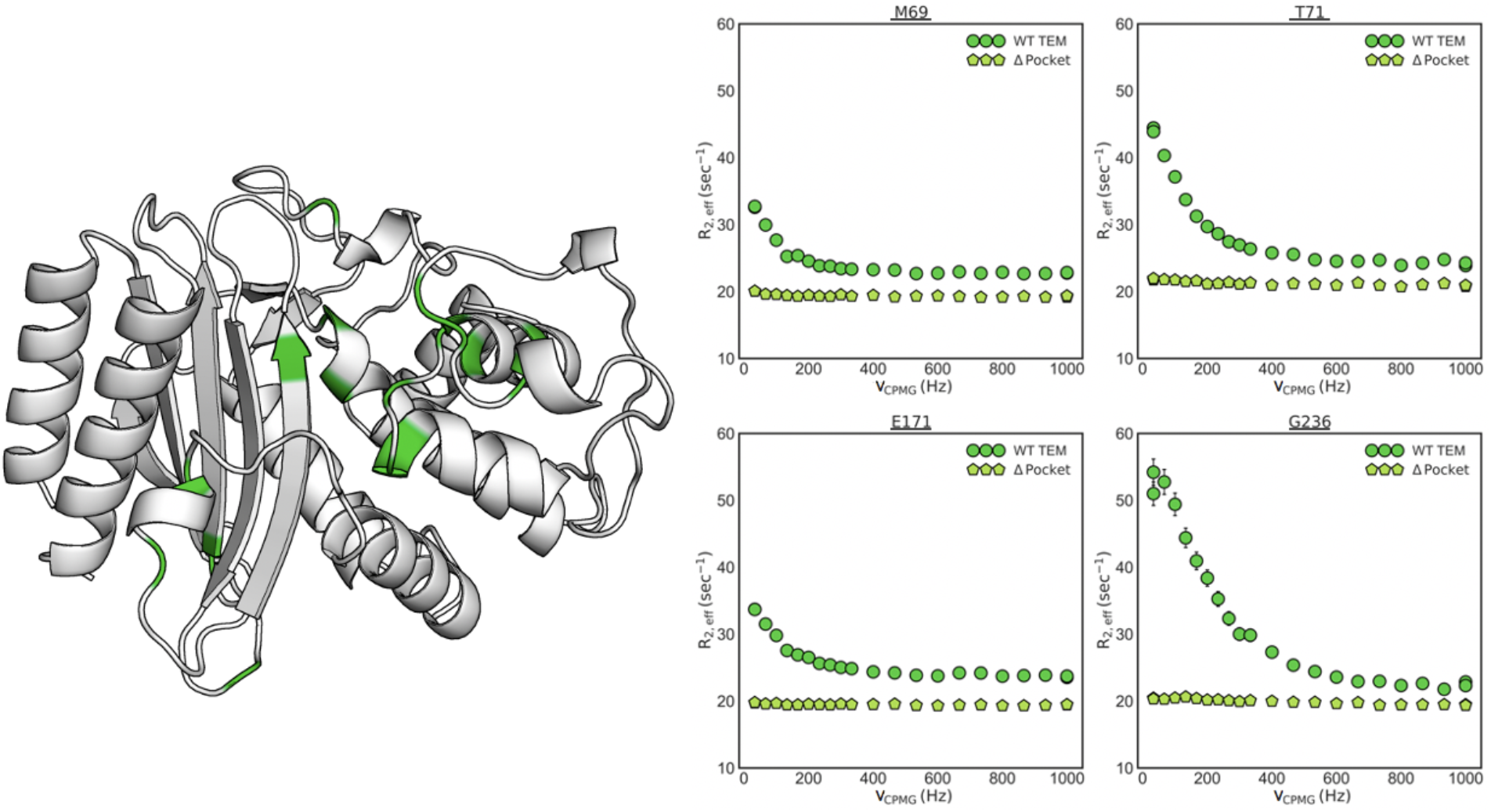
The R241P mutation in TEM removes dynamics on the microsecond to millisecond timescale as monitored by relaxation dispersion. (Left) Highlighted in green on the wild type (WT) TEM structure (PDB: 1xpb) are the residues showing conformational exchange as established by Car-Purcell-Meiboom-Gill (CPMG) experiments. (Right) ^15^N CPMG profiles for wild type TEM (green circles) and TEM R241P, a variant with no Ω-loop pocket, (light green pentagons) are shown for a set of representative residues.

**Supplemental Figure 7.**
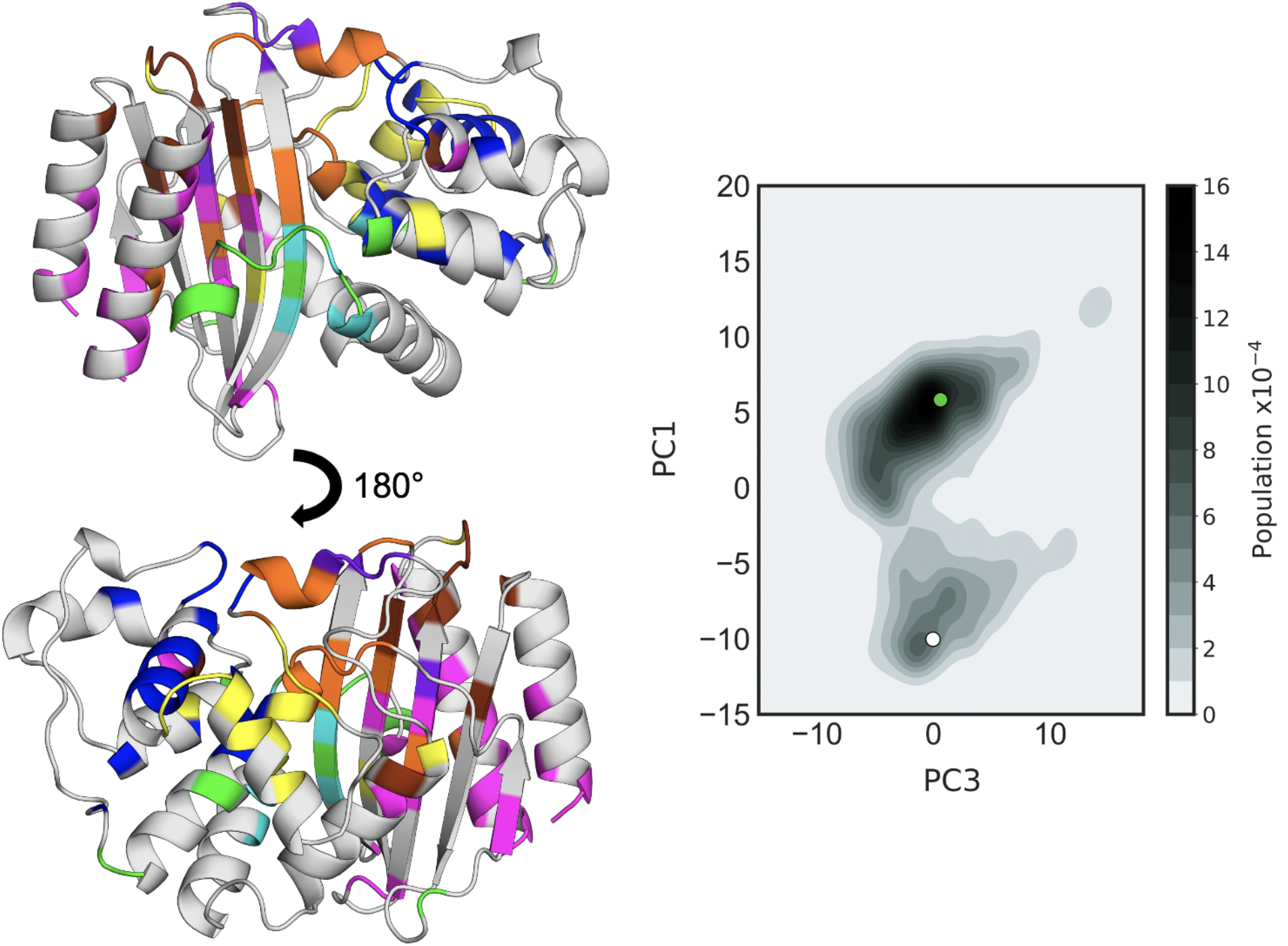
Simulation data was analyzed using CARDS and PCA. (Left) Each color highlighted on the WT TEM structure (pdb: 1xpb) represents a different CARDS community. These communities are residues which have correlated dihedral motions. The orange community includes the Ω-loop and the catalytic S70. (Right) We performed PCA on the orange community and found two resolved minima. The colored circles represent the positions of the exemplar structures shown in main text figure 4b.

**Supplemental Figure 8.**
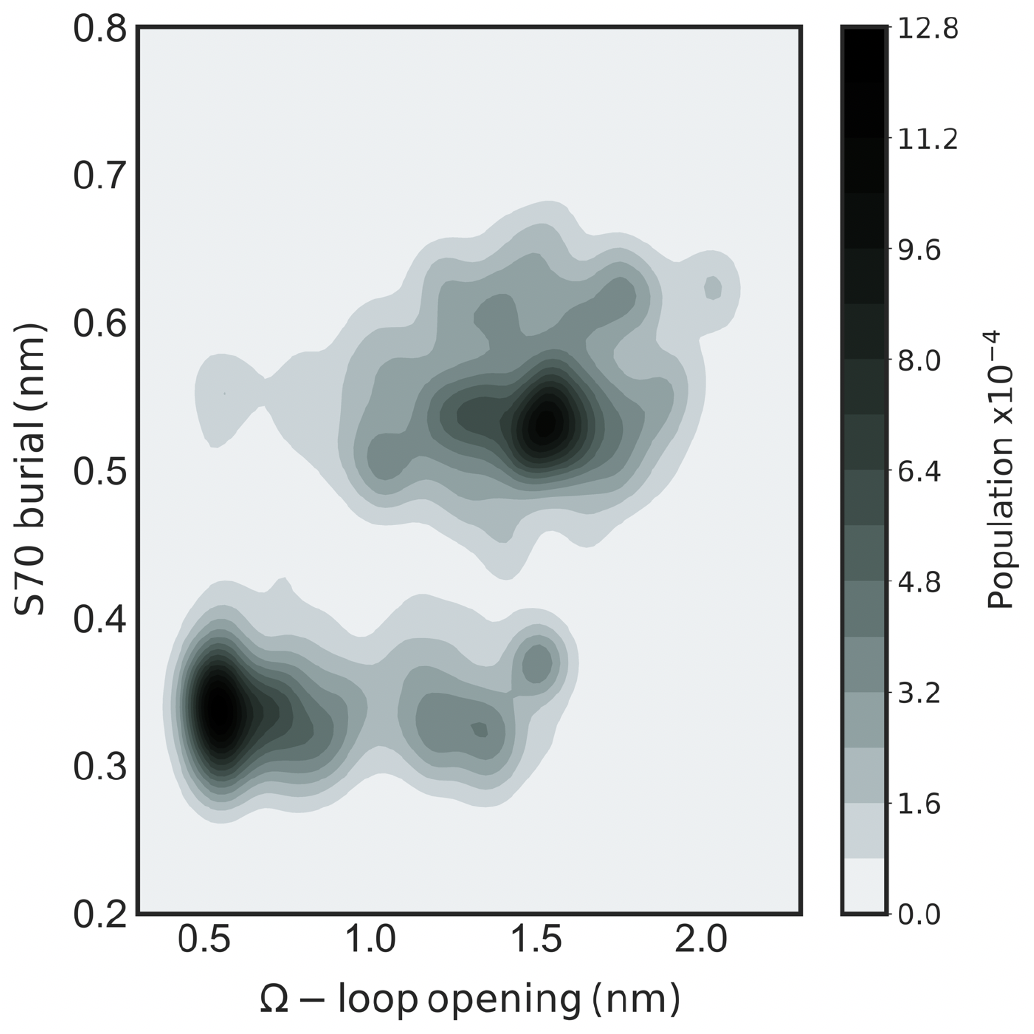
Ω-loop pocket opening is correlated with S70 burial. A 2D histogram of catalytic S70 burial as a function of Ω-loop pocket opening shows that when the pocket is open, S70 is predominantly buried. S70 burial is captured using a backbone hydrogen bond distance between S70 and K73. The Ω-loop opening distance is characterized by the Cα-Cα distance between E240 and E171.

**Supplemental Table 1.** In order to determine existence of the Ω-loop pocket, the pocket population should be higher than the unfolded population.

| TEM Variant | Ω-Loop Pocket Open Population (%) | Unfolded Population (%) |
| --- | --- | --- |
| Wild Type (S243C) | $1.1 \pm 0.2^\dagger$ | $0.02 \pm 0.01^\dagger$ |
| E240D/S423C | $4 \pm 1^\ddagger$ | $0.0047 \pm 0.0009$ |
| R241P/S243C | $0.002 \pm 0.005^\diamond$ | $0.12 \pm 0.07$ |
| E240D/R241P/S243C | $0.19 \pm 0.01$ | $0.14 \pm 0.02$ |

$^\dagger$ Porter *et al.*, 2019
$^\ddagger$ Labeling in the EXX regime, individual $k_{op}$ and $k_{cl}$ were determined and then divided to obtain the $K_{op/cl}$
$^\diamond$ Determined to not have an Ω-loop pocket as the pocket population is less than the unfolded population

**Supplemental Figure 9.**
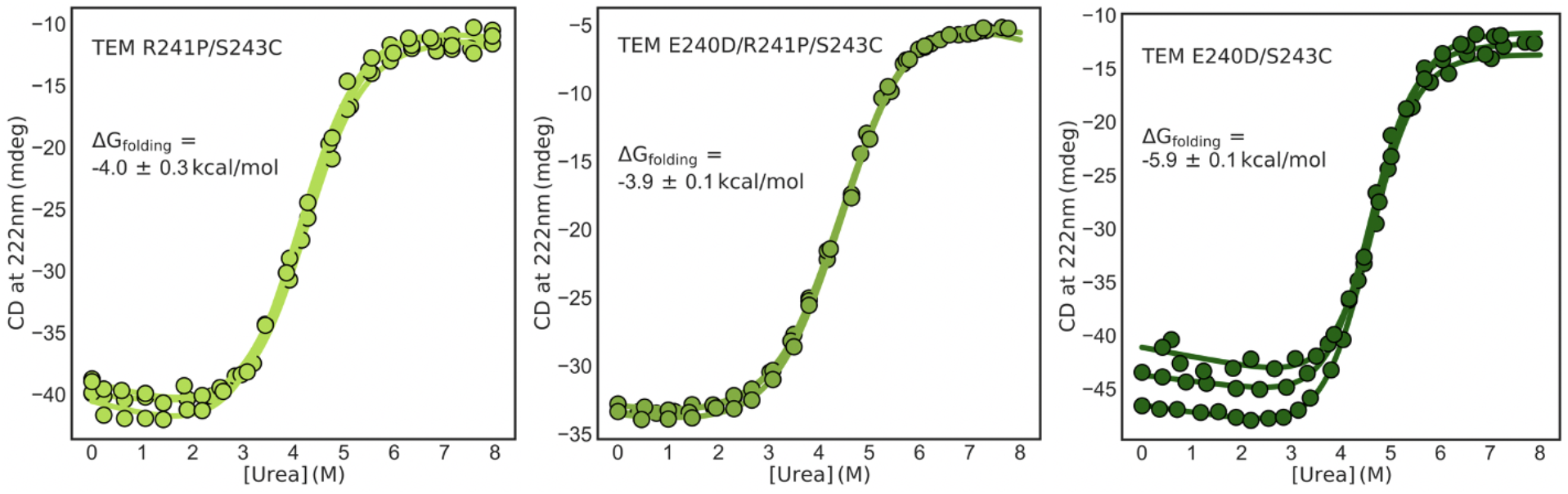
The labeling rates for TEM R241P are on the order of magnitude as those expected for labeling due to protein unfolding, while the rates for TEM E240D are much faster. Urea denaturation was followed by circular dichroism for TEM R241P (left), TEM E240D/R241P (middle), and TEM E240D (right). Error is reported as the standard deviation of three replicate experiments.

**Supplemental Figure 10.**
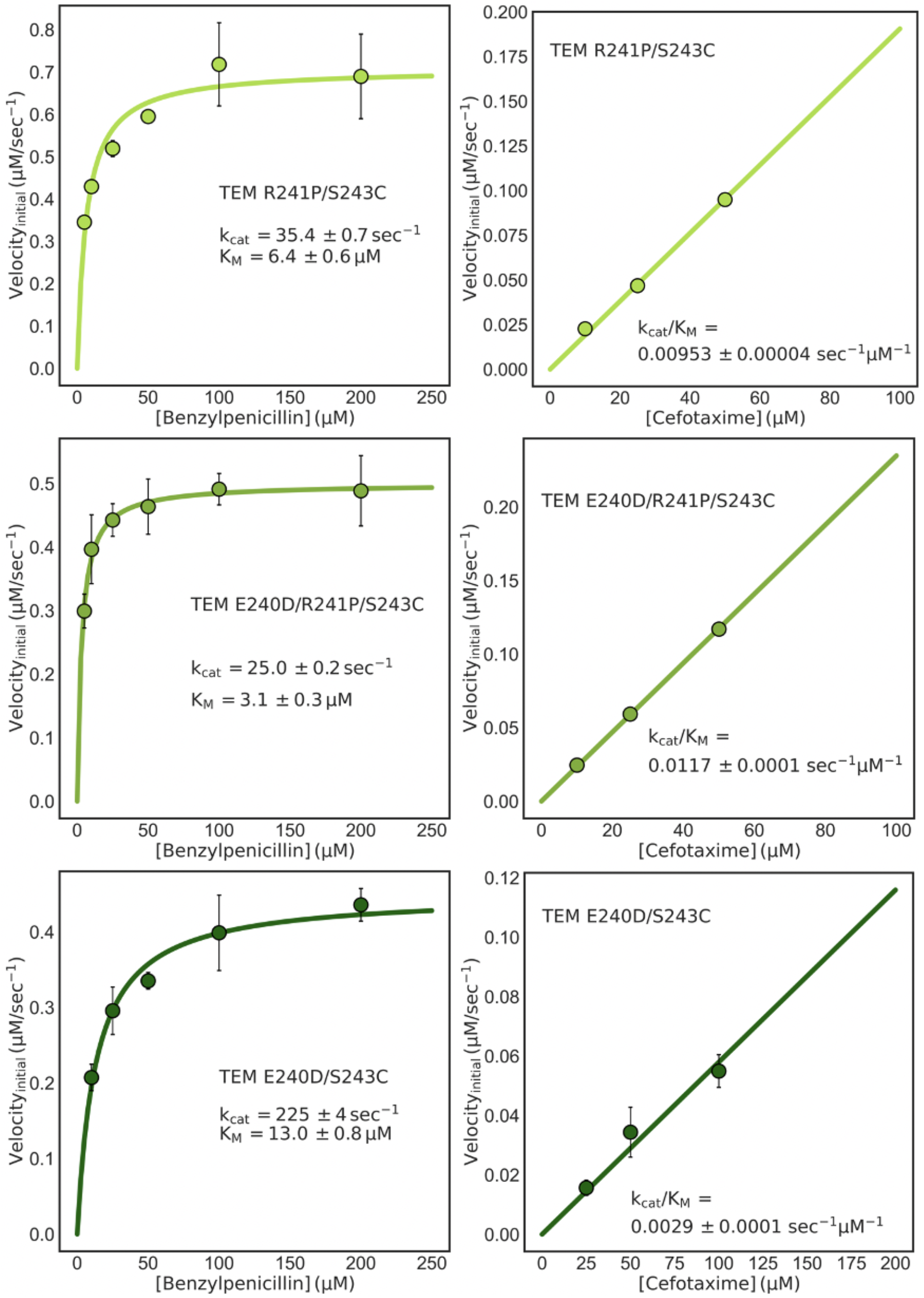
The R241P mutation increases cefotaxime activity and decreases benzylpenicillin activity, while the E240D mutation decreases cefotaxime activity and increases benzylpenicillin activity. The full Michaelis-Menten equation was fit to the benzylpenicillin activities. A line with a slope equal to the catalytic efficiency multiplied by the enzyme concentration was fit to the cefotaxime activities. Error bars are shown as the standard deviation of three replicate measurements. Error for each parameter was determined using bootstrapping.

### Supplemental Methods

#### NMR CPMG experiments

We recorded ^15^N CPMG experiments as previously described,[54] using a constant-time relaxation interval, T_relax_, of 30 milliseconds.[55] We sampled 20 V_CPMG_ values of ≤1 kHz, using CPMG refocusing pulses applied at a γB_1_/2π = 6 kHz field and phase-modulated according to the {x,x,y,-y} cycling scheme.[56] We applied a γB_1_/2π = 15.6 kHz field ^1^H continuous wave decoupling during T_relax_. To ensure constant heating in the reference experiments (recorded with T_relax_ equal to 0 seconds), we applied the same ^1^H continuous wave decoupling immediately prior to the recycle delay.

#### Urea denaturation experiments

We prepared samples of 35 μg/mL protein in 50 mM potassium phosphate, pH 7.0 at various concentrations of urea and equilibrated the samples at room temperature overnight. After one-minute incubation in an Applied Photophysics Chirascan equipped with a Quantum Northwest Inc. TC125 Peltier-controlled cuvette holder, we monitored circular dichroism (*CD*) signal at 222 nm. We recorded the signal for one minute at 25°C in a one cm path length quartz cuvette. Then, we measured the refractive indexes of each sample in order to the determine their precise urea concentration. We determined the free energy values by fitting a two-state folding model (below) to the CD data and using the linear extrapolation method.[57] Each variant was measured in triplicate experiments.

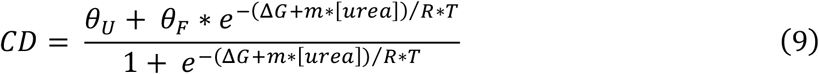

Here, *θ_U_* and *θ_F_* are the CD signals for the unfolded and folded states, fit as lines. Δ*G* is the extrapolated free energy difference between the unfolded and folded states in the absence of urea, and *m* is the proportionality constant related to the steepness of the folding transition. *R* is the gas constant and *T* is temperature.

#### Urea unfolding kinetics

We prepared a protein sample in 50 mM potassium phosphate, pH 7.0 and samples of various urea concentrations above the concentration midpoint (*C_M_*) in the same buffer. After five-minute incubation in an Applied Photophysics Chirascan equipped with a Quantum Northwest Inc. TC125 Peltier-controlled cuvette holder, we added the protein to the urea buffer, diluting the protein to a final concentration of 35 μg/mL, and manually mixed by inverting the cuvette. We then monitored the circular dichroism (*CD*) signal at 222 nm over time at 25°C in a one cm path length quartz cuvette. We measured the refractive indexes of each sample in order to the determine their precise urea concentration. An exponential was fit to each unfolding kinetic trace to determine the observed unfolding rate at that given urea concentration. The unfolding rate in the absence of urea was then calculated using a linear extrapolation fit to the log of the observed unfolding rates as a function of urea.

